# Loss of MeCP2 leads to sleep deficits that are time-of-day dependent and worsen with sleep deprivation

**DOI:** 10.1101/2025.04.16.649209

**Authors:** Abrar Al Maghribi, Michael Rempe, Elizabeth Medina, Kaitlyn Ford, Kristan Singletary, Caitlin Ottaway, Lucia Peixoto

## Abstract

Rett syndrome (RTT) is a severe, progressive neurodevelopmental disorder caused by mutations in the X-linked gene encoding methyl-CpG-binding protein 2 (MECP2). Sleep problems are frequently reported in Rett Syndrome, but the exact nature remains relatively unexplored. Currently there is limited understanding the role of MECP2 in sleep architecture and regulation. In this study, we employed longitudinal electroencephalographic (EEG) and electromyographic (EMG) recordings to investigate sleep architecture during baseline conditions as well as the homeostatic response to sleep deprivation (SD) in Mecp2-/y male mice. At baseline, Mecp2-/y mice have more non-rapid-eye-movement (NREM) sleep and less rapid-eye-movement (REM) sleep than their wildtype littermates during the light period. However, Mecp2-/y mice display altered sleep timing during the dark period, spending more time in both NREM and REM during the first half and less time during the second half. We also observe differences in REM sleep and wake quality based on spectral properties of the EEG. In response to SD, Mecp2-/y mice can accumulate and discharge sleep pressure normally and show a sleep rebound. However, baseline differences in sleep architecture are heightened after SD. Overall, our findings show that RTT mice exhibit distinct sleep patterns compared to wildtype mice, with time-of-day-dependent variations in NREM and REM sleep, as well as altered EEG spectral properties, that become more pronounced following SD. Future research should explore the molecular mechanisms through which MECP2 regulates circadian sleep architecture to develop targeted therapeutics for sleep disturbances in RTT patients.

**Highlights:**

- Mecp2^−/y^ mice show time-of-day-dependent alterations in NREM and REM sleep.
- EEG analysis revealed distinct sleep and wake quality in Mecp2^−/y^ mice.
- Sleep deprivation exacerbates baseline sleep architecture differences.
- Longitudinal EEG/EMG recordings captured comprehensive sleep patterns.

## 1. Introduction

Rett syndrome (RTT) is a rare neurodevelopmental disorder that affects 1 in 10,000– 15,000 females (Chahrour & Zoghbi, 2007). The majority of classic RTT cases (<95%) are caused by mutations in the X-linked gene encoding methyl-CpG–binding protein 2 (*MECP2*) (Lombardi et al., 2015; Zoghbi, 2005). Clinically, RTT is associated with a wide range of neurological symptoms, including delayed growth, motor impairments such as repetitive hand-wringing, cognitive delays, communication challenges, autistic features, perinatal death (Amir et al., 1999; Hagberg et al., 1983; Neul, 2012) and sleep problems (Raspa et al., 2024). Sleep disturbances, including difficulty falling asleep, frequent night awakenings, and disrupted sleep patterns, are prevalent in this population. These disruptions often start during early childhood and persist throughout life (Glaze et al., 1987; Zhang et al., 2023). These sleep issues not only impact the quality of life of those with RTT, but also place additional strain on caregivers, highlighting the importance of addressing sleep as a key component in the overall management of the disorder. Thus, understanding the underlying mechanisms and providing tailored interventions can be crucial in improving sleep quality and, by extension, the well-being of individuals with RTT and their families. Translational models can provide insight into the mechanisms underlying sleep problems in RTT.

Although Rett syndrome primarily affects females, female *Mecp2*-mutant mice are not always ideal models due to the effects of X-chromosome inactivation (XCI). Because females have two X-chromosomes, one is randomly silenced in each cell, leading to a mosaic pattern where some cells express the mutant *Mecp2* gene and others do not. This cellular variability makes it difficult to recognize which phenotypes result from cell-autonomous effects or from interactions with neighboring cells (non-autonomous effects). Therefore, many studies use hemizygous *Mecp2*-mutant male mice which have a total absence of the MeCP2 protein and they display more consistent phenotypes. (Vashi & Justice, 2019). However, there are limited studies regarding how the loss of *MECP2* gene affects sleep. There is a scarcity of studies looking at EEG in *Mecp2*-null male mice lacking the MeCP2 protein. Only one study has studied sleep in this population, and it reported no significant difference in total sleep time between mutants and wildtype mice (Johnston et al., 2014a). However, that study did report that mutant mice show an altered distribution of sleep during the dark period, suggesting a time-of-day-dependent effect. Additionally, the study proposes that the mutation impacts slow-wave delta power. However, no study has evaluated the homeostatic response to SD. In this study we carried out a full characterization of sleep architecture and regulation using the gold-standard combination of electroencephalography (EEG) and electromyography (EMG). Overall, our findings indicate that the absence of MECP2 has a significant impact on sleep, which is dependent on the time of day and exacerbated by loss of sleep.

## 2. Material and methods

### 2.1 Experimental model and subject details

Experimental subjects were obtained by breeding heterozygous female B6.129P2(C)- Mecp2^tm1.1Bird^/J to wildtype (WT) male 129S6SvEvTac mice to produce *Mecp2*^*-/y*^ and WT male littermates (n = 10 WT, 10 *Mecp2*^*-/y*^). Animals were housed in a temperature and humidity-controlled environment under a 12-hour light/dark cycle and given access to food and water *ad libitum*. All experimental procedures were approved by the Institutional Care and Use Committee of Washington State University and in accordance with the National Research Council guidelines and regulations for experiments in live animals.

### 2.2 Surgical procedures

*Mecp2*^−/y^ and WT littermates control mice between the ages of 6 and 8 weeks old underwent stereotaxic implantation with electroencephalographic (EEG) and electromyographic (EMG) electrodes as described in our previous work (Ingiosi et al., 2019; Medina et al., 2022). Briefly, four stainless steel screws (BC-002MPU188, Bellcan International Corp, Hialeah, FL) were implanted into the skull above the frontal (2) and parietal (2) cortices, and 2 EMG electrodes were inserted into neck muscles for EMG recording.

### 2.3 EEG/EMG data acquisition and processing

Mice were individually housed in cylindrical polypropylene containers tethered with lightweight EEG/EMG cables in sound attenuated enclosures (sleep chambers). Mice were given a minimum of 5 days to acclimate to the tethering and recording environment prior to recording. EEG/EMG recordings consisted of 24 hours of baseline (BL) starting at lights on (ZT=0) under undisturbed conditions followed by 5 hours of SD and 19 hours of subsequent recovery sleep as in our previous work (Ingiosi et al., 2019; Medina et al., 2022). A soft brush was used to gently stroke the animals when needed to ensure constant wakefulness during the 5 hours of SD. Sleep deprivation efficiency was measured by calculating the percentage of wake epochs over the 5-hour SD period. Both WT and *Mecp2*^−/y^ mice exhibited SD efficiency above 95%, confirming that the animals remained awake throughout the deprivation period.

An RHD2000 amplifier Intan system (16-channel RHD USB Recording System, Intan Technologies, Los Angeles CA) was used to record the EEG and EMG signals. The amplifier rate was set to 1 kilo-sample per second with a cutoff at 0.01 Hz, lower bandwidths at 0.1 Hz, upper bandwidth at 200 Hz, and notch filtered at 60 Hz. Raw data was collected in (.rhd) format and then converted to European Data Format (.edf) via custom software. Vigilance states were initially scored in 4-second epochs using the deep-neuronal network algorithm SPINDLE (Miladinović et al., 2019) and then verified manually using SleepSign software (Kissei Comtec CO., LTD). Data was scored into either wake, non-rapid eye movement sleep (NREM) or rapid eye movement sleep (REM). State scoring and data analysis were conducted in a blinded and randomized manner. A single file containing all data, scored states, as well fast Fourier transform (FFT) data was exported from SleepSign and used for subsequent analyses. EEG epochs with artifacts were omitted from spectral analysis. EEG spectra power was obtained from 0-50 Hz with 0.244 Hz bins and normalized relative to total power across states and presented in smooth curved lines with 95% confidence intervals displayed around each spectrum. The total recording time (TRT) was presented as the percentage of the total state-specific time during the light and dark periods. Sleep pressure (normalized delta power) was defined as power in the delta range (0.5-4 Hz) of NREM during recovery sleep and normalized relative to the baseline NREM delta power during the last 4 hours of the light period. Sleep onset latency was defined as the time elapsed from the end of SD to the first episode of NREM lasting at least 28 seconds. To identify differences in sleep fragmentation, bout number and time were also analyzed. A bout was defined as a continuous 2 number of epochs in each state.

### 2.4 Statistical analysis

Data analysis was performed using MATLAB (MathWorks). Hourly time-in-state data are presented as means ± standard error of the mean (SEM). Repeated measures ANOVAs were used to determine significant differences between RTT mice and WT mice for hourly time spent in each arousal state, and normalized NREM delta power after SD. Sleep latency as well as 12-hour average values for bout number and bout duration used ANOVAs. To compare the spectral content of the EEG between RTT mice and WT mice, we plotted normalized spectral power and visually inspected for overlap of 95% confidence intervals. In case of a significant main effect of genotype or interaction between genotype and time, t-tests were used in post-hoc analyses and corrected for multiple comparisons using the Benjamini Hochberg procedure. p-values <0.05 were considered statistically significant.

## 3. Results

### 3.1 Absence of MECP2 influences sleep in a time-of-day dependent manner

The goal of this study was to investigate whether deletion of the MECP2 gene produced deficits in sleep or the homeostatic response to acute SD. To do so, we performed EEG/EMG recordings under baseline conditions for 24 hours, followed by 5 hours of SD and 19 hours of subsequent recovery sleep. We first investigated whether there were differences in sleep quantity during both the light period (when mice sleep the most, rest phase) and the dark period (when mice sleep the least, active phase) under baseline conditions. Figure 1 shows the time spent in each arousal state (Wake, NREM, and REM sleep) across baseline measurements in the light period and the dark period. During the light period, *Mecp2*^−/y^ mice do not have a significant difference in wake time, however a significant difference in time in NREM and a significant decrease in REM is present. During the dark period we detected a significant interaction between genotype and time. *Mecp2*^−/y^ mice spend more time in wake during the second half of the dark period. In addition, we observed an increase in NREM during the first half of the dark period followed by a decrease during the second half, a pattern that is also seen during REM sleep. To determine if the changes in total time in state were caused by a sleep fragmentation, we evaluated state specific bout numbers and bout durations. *Mecp2*^−/y^ mice show a reduction in number of REM bouts in the light period and the second half of the dark period (Supplementary Material, Figure 1A). REM bout duration was also reduced in *Mecp2*^−/y^ mice compared to WTs during the light period. Interestingly, bout duration was reduced for all states in the first half of the dark period. (Supplementary Material, Figure 1B). Full statistics for Figure 1 are available in Additional File 1 (Sheet: Figure 1. Time in State.BL). Bout analysis statistics can be found in the sheet titled (Supplementary Figure 1. Bout BL).

**Figure 1.**
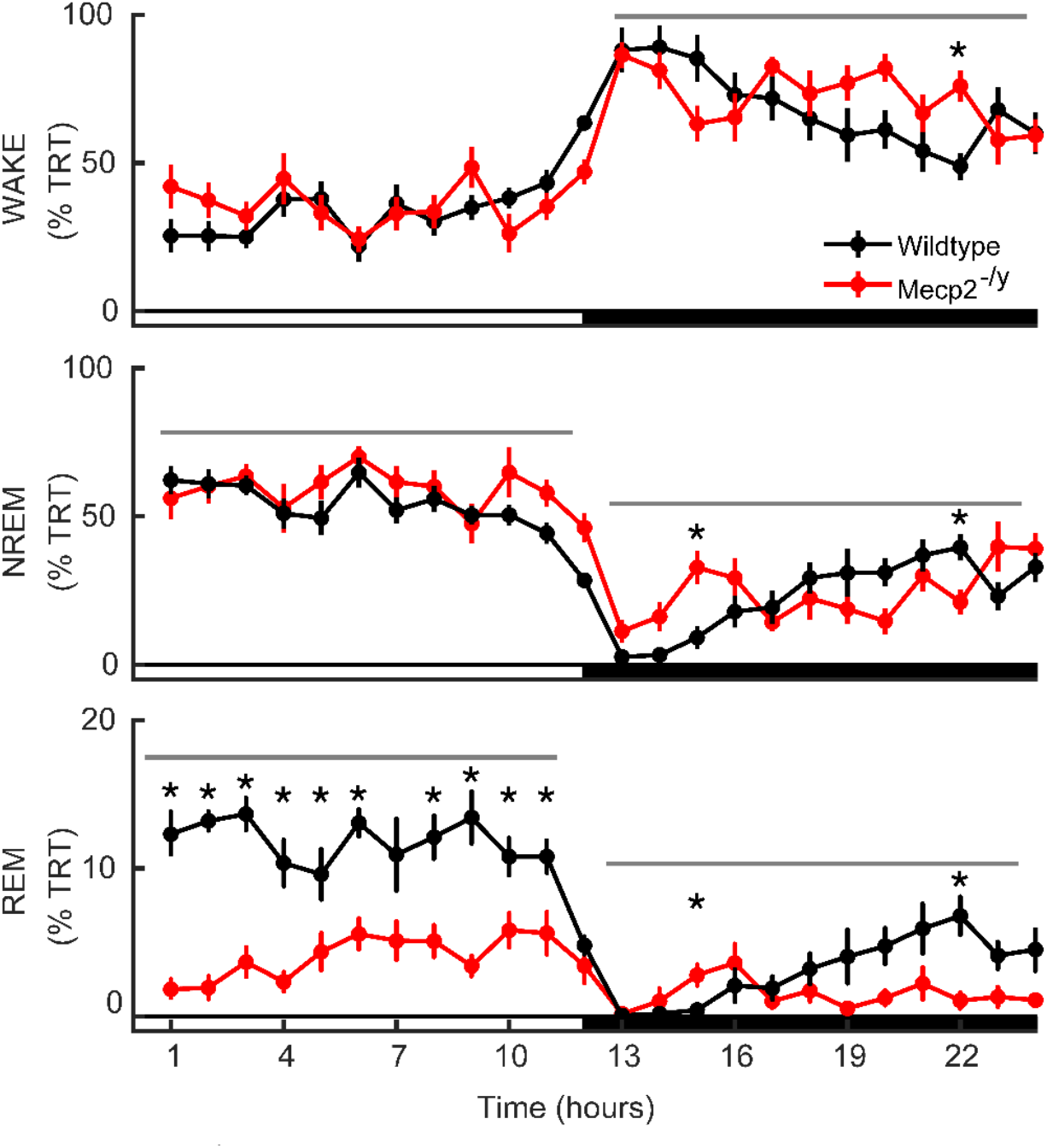
*Mecp2*^−/y^ mice sleep differently during the light and dark period. Time in state is displayed as percentage total recording time (TRT) during baseline for wake (top row), NREM (middle row) and REM (bottom row). The light period (white-filled bar on x-axis) hours 0 to 12 and dark period (black-filled bar on x-axis) hours 13 to 24 tested separately. Significant measurements were determined using repeated measures-ANOVA represented with gray bar with post hoc pairwise comparisons using Benjamini Hochberg correction; *p<0.05 to compare the main effect of genotype over 12 hours. WT represented in black (n=10), Mecp2^−/y^ represented in red (n=10).

Overall, the absence of MECP2 produces significant sleep deficits that are dependent on time of day. During the light period we observe a reduction of REM sleep amount, bout number and duration. Genotype differences during the dark period vary depending on time of day and display a paradoxical effect in which RTT mice sleep more when mice are usually most active (first half of the dark period) and less when they usually take a nap (second half of the dark period).

### 3.2 *Mecp2* mutant mice display differences in spectral quality in both Wake and REM

To better understand qualitative differences in sleep and wake architecture, we conducted spectral analysis- a quantitative method that examines the frequency structure of brain activity (Figure 2). This analysis provides insights into frequency-specific components of brain activity offering a deeper understanding of the physiological changes associated with MECP2 deficiency. *Mecp2*^−/y^ mice show a left shifted distribution of wake spectra, in which we observe lower power in the 7-15 Hz range compared to WT (Figure 2A). The same phenomenon can be observed in NREM sleep (Figure 2B). In addition, we also observe an increase in power at lower frequencies in REM (Figure 2C). We observe the same spectral differences pattern in both the light and dark periods, despite the differences in sleep amounts (Supplementary Material, Figure 2). Overall, the absence of MECP2 appears to be associated with a decrease in power during wake, and NREM and an increase in lower frequencies during REM.

**Figure 2.**
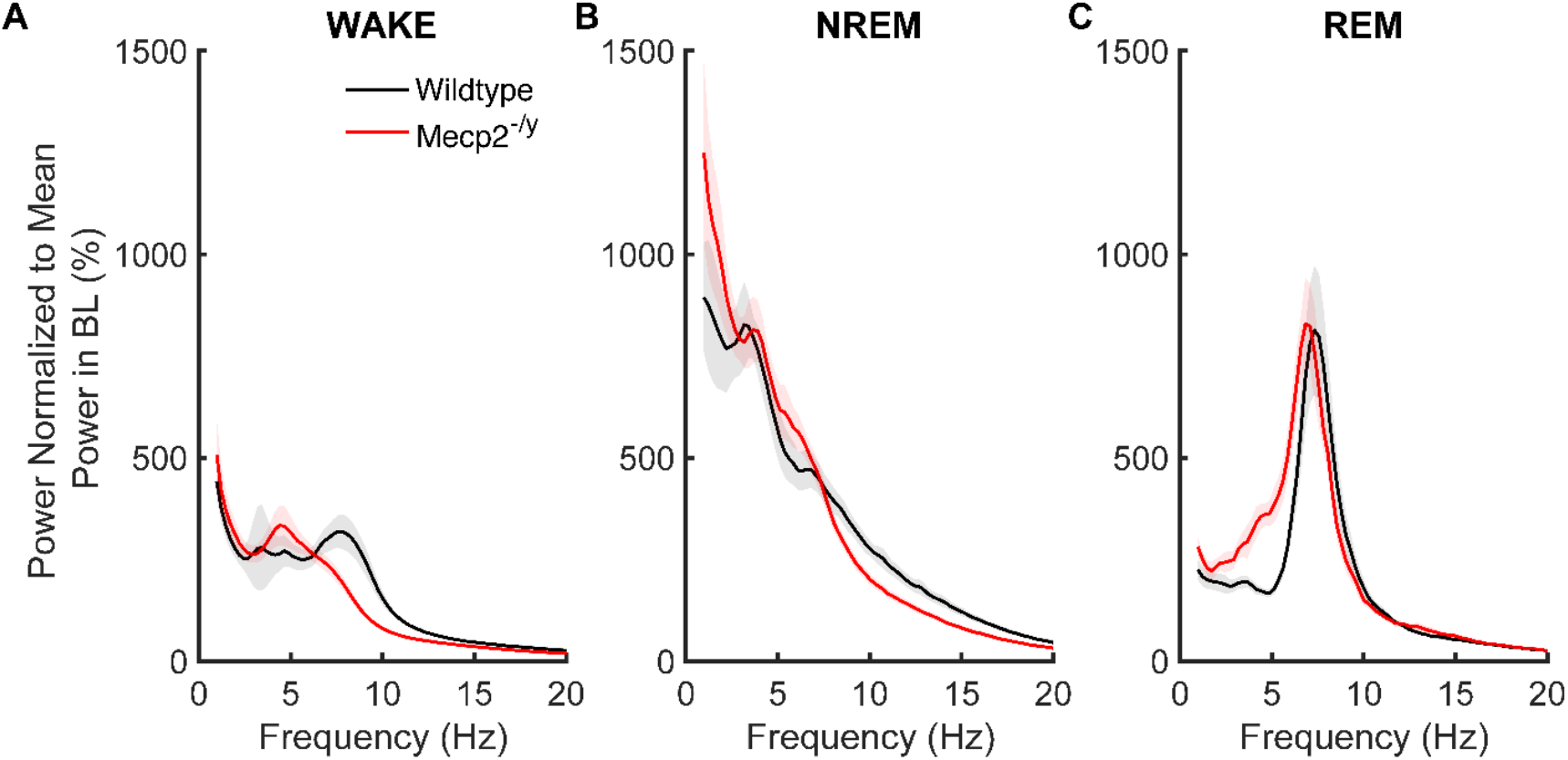
*Mecp2*^−/y^ mice show altered frequency for baseline spectra. Normalized EEG spectra power in (A) wake, (B) NREM, and (C) REM at baseline. In each case power was normalized to the mean power in all EEG frequencies averaged across all sleep states. The red line represents *Mecp2*^−/y^, and the black line represents the wild type. 95% confidence intervals are displayed around each spectrum, light gray for WT, and light red for *Mecp2*^−/y^. WT is represented in black (n=10), *Mecp2*^−/y^ is represented in red (n=10). Baseline (BL).

### 3.3 Absence of MECP2 does not inhibit the homeostatic response to sleep deprivation

The homeostatic regulation of sleep is a fundamental process that is used to ensure the brain compensates for sleep deprivation. After a period of acute sleep deprivation, animals respond by falling asleep faster and sleeping longer and deeper. The standard metric of sleep pressure is the increase in power in the delta frequency range (1-4 Hz.) of NREM sleep after SD, relative to baseline (normalized delta power) (Borbely, 1982). *Mecp2*^−/y^ mice show an increase in NREM delta power reflecting a proper homeostatic response to sleep loss (Figure 3A). In addition, quantification of the latency to sleep following SD revealed no significant genotype difference in sleep latency (Figure 3B). Lastly, both WT and *Mecp2*^−/y^ mutants increased sleep amounts after SD compared to baseline (Figure C). Overall, these findings indicate that MECP2 does not inhibit the homeostatic response to SD. Full statistics for Figure 3 can be found in Additional File 1 ((Sheet: Figure 3).

**Figure 3.**
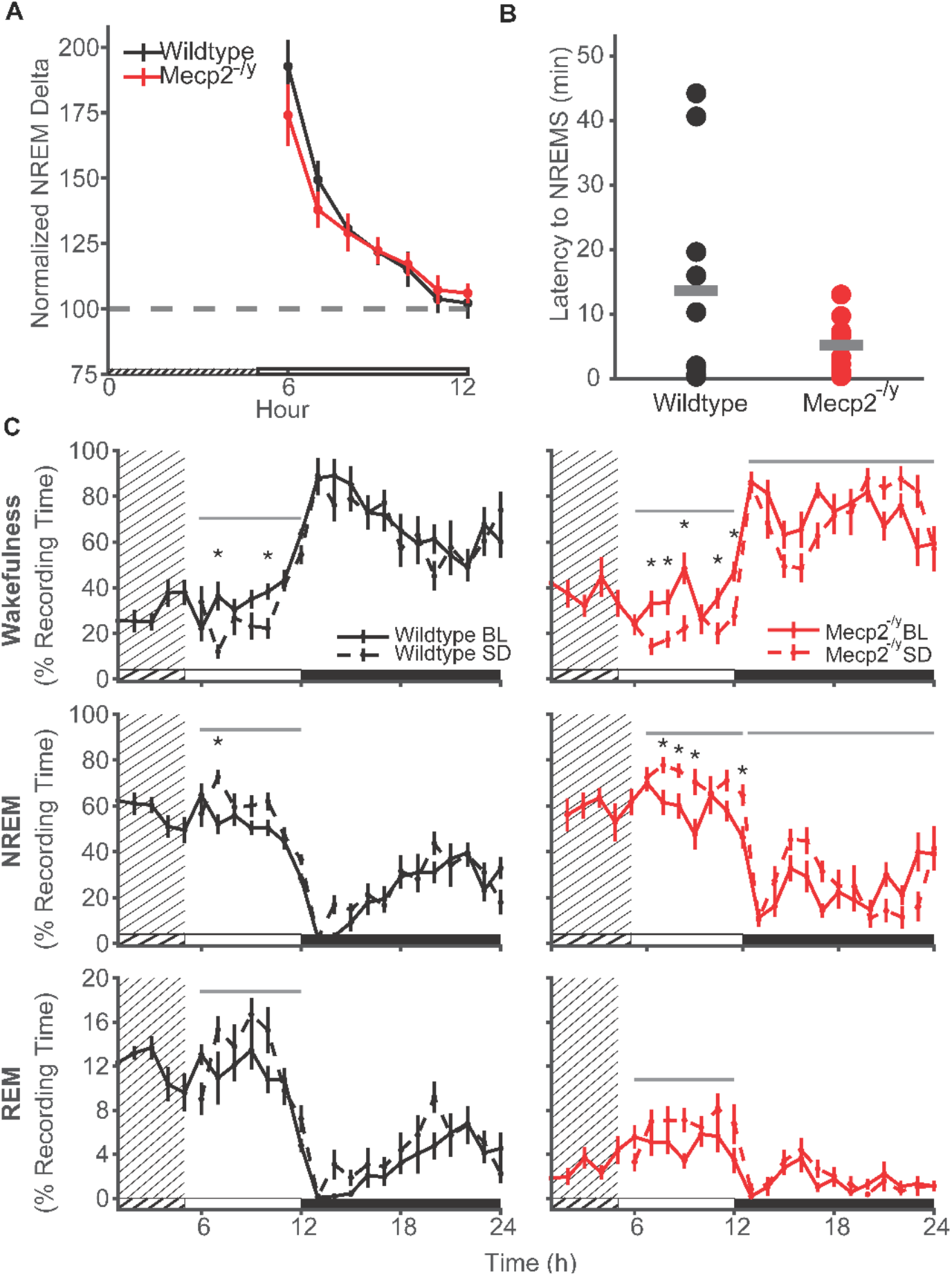
*Mecp2*^−/y^ mice show a typical homeostatic response post sleep deprivation. (A) Hourly NREM sleep delta power (0.5-4 Hz) during recovery sleep post sleep deprivation. Differences between genotypes were tested using repeated measures-ANOVA. Error bars represent SEM, grated area represents the five hours of SD. (B) Sleep latency to NREM sleep after sleep deprivation. Data were analyzed using an unpaired t-test. (C) The percentage of total recording time (TRT) spent in Wake (top row), NREM sleep (middle row) and REM sleep (bottom row) in recovery sleep compared to baseline. The gray bar above plots represents a significant difference between the two curves as indicated by a repeated measures ANOVA across hours 6-12 (light period) or 13-24 (dark period). The light period is represented by a white bar, and the dark period black bar along the bottom of each panel. Dashed lines represent data from the recovery sleep day, while solid lines represent data from the baseline day, grated area represents the five hours of SD. WT is represented in black (n=10),and *Mecp2*-/y is represented in red (n=10).

### 3.4 Sleep deprivation exacerbated the baseline time in state differences of Mecp2^−/y^ mice

Finally, we examined the total time spent in wake, NREM, and REM sleep as well as spectral analysis of recovery sleep. Figure 4 illustrates the total time spent in each state (wake, NREM, and REM sleep) across the 24-hour period. During the light period we detect a decrease in wake, an increase in NREM sleep, and a significant decrease in REM in Mecp2^−/y^ sleep. During the dark period we detect a significant interaction between genotype and time, which is similar to baseline sleep. *Mecp2*^−/y^ mice spend significantly less time in wake during the first half of the dark period and more time in wake during the second half of the dark period. Moreover, *Mecp2*^−/y^ mice spend more time in NREM during the first half of the dark period followed by a decrease during the second half, a pattern that is also seen during REM sleep. Bout analysis revealed that *Mecp2*^−/y^ mice exhibited a significant increase in NREM bout number during the first half of the dark period and a significant reduction of REM bout number during the second half of the dark period (Supplementary Material, Figure 3A). However, *Mecp2*^−/y^ mice show a significant reduction in wake bout duration during the light period and the first half of the dark period and a significant reduction in NREM bout duration during the second half of the dark period (Supplementary Material, Figure 3B). Interestingly, REM bout duration was reduced for *Mecp2*^−/y^ mice for both light and both halves of the dark period (Supplementary Material, Figure 3B). Our findings indicate that sleep deprivation exacerbates the baseline phenotypic characteristics of *Mecp2*^−/y^ mice. Spectral analysis of recovery sleep (Supplementary Material, Figure 4) shows the same deficits seen at baseline (Figure 2). Full statistics for recovery sleep analysis can be found in Additional File 1 (Sheet: Figure 4. Time in State.SD). Bout analysis statistics can be found in the Additional File 1 sheet titled (Supplementary Figure 3. Bout SD).

**Figure 4.**
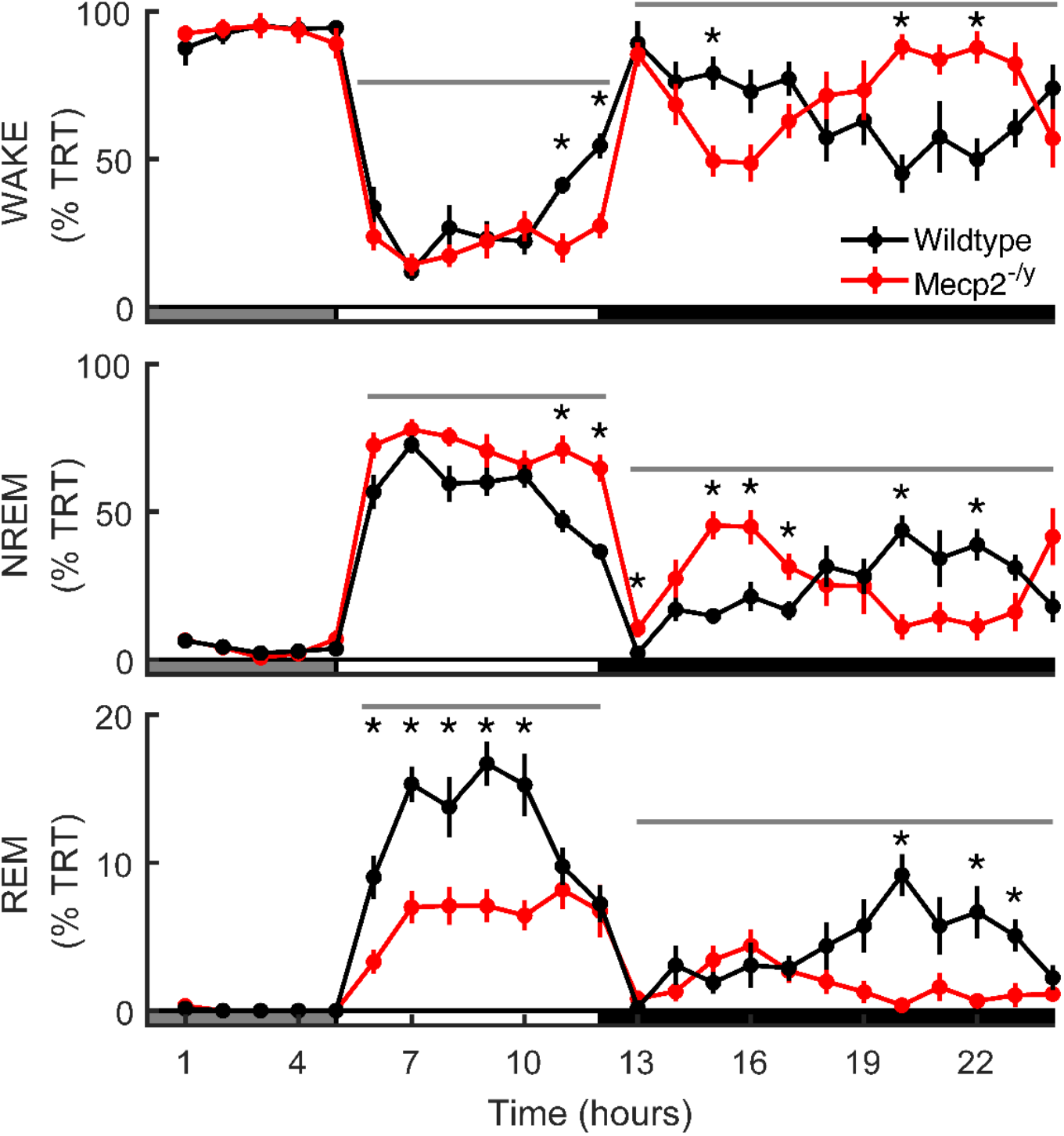
Sleep deprivation exacerbates the baseline phenotypic characteristics of *Mecp2*-/y mice. Time in state is displayed as the percentage total recording time (TRT) after sleep deprivation for percentage of each hour spent in (A) wake, (B) NREM, or (C) REM. The light period is shown as a white bar on the horizontal axis and the dark period is shown with a black bar. The grated area represents the five hours of SD. Significant differences between genotypes over 12 hour or 6 hour segments are represented with a gray bar and determined using repeated measures-ANOVA. Asterisks indicate specific time points where post-hoc tests showed significant differences. (All post-hocs were corrected using Benjamini-Hochberg). WT represented in black (n=10), *Mecp2*-/y represented in red (n=10).

## 4. Discussion

RTT is caused by the loss-of-function mutations in *MECP2*. While MECP2 is known to be crucial for normal brain development, its role in sleep regulation remains unclear. In this study, we used the *Mecp2*-null mouse model to characterize the *Mecp2*^−/y^ mice’s sleep architecture during baseline and in response to sleep deprivation.

At baseline, our findings show that *Mecp2*^−/y^ mice have significantly more NREM sleep and significantly less REM sleep compared to WT mice during the light period. However, during the second half of the dark period, *Mecp2*^−/y^ mice spend significantly less time in NREM and REM (Figure 1). These findings indicate that the absence of MECP2 affects sleep differently across the light and the dark periods and it is thus dependent on the time of day. Changes during the dark period may indicate dysregulation in the transition between vigilance states. For instance, *Mecp2*^−/y^ mice sleep more in the first half of the dark period (when mice are usually awake) and less in the second half of the dark period (when mice usually take a nap). Interestingly, RTT patients also report having disrupted sleep-wake cycles (Piazza et al., 1990; Zhang et al., 2023). In addition, our study demonstrates that *Mecp2*^−/y^ mice display a strong REM sleep phenotype. *Mecp2*^−/y^ mice display reduced REM sleep amount, bout number, and bout duration most of the day, as well as altered spectral frequency in REM sleep. This is in agreement with clinical studies reporting a reduction in REM sleep in RTT patients (Amaddeo et al., 2019; Ammanuel et al., 2015; Glaze et al., 1987). A previous *Mecp2*^−/y^ mice study reported no significant differences in overall wake and sleep time percentages between WT and mutant mice (Johnston et al., 2014) However, when activity was analyzed separately by light and dark phases, mutants exhibited more consolidated wake bouts, characterized by longer wake cycle durations, fewer bouts, and a significantly longer duration of the longest wake cycle. Additionally, the study found that mutant mice had a significantly longer duration of the longest NREM episode compared to WT (Johnston et al., 2014). In contrast, we only observe differences in REM sleep structure (bout number and duration) but not in NREM. Although the homeostatic response to SD is intact, SD amplified the baseline sleep phenotype. This indicates that *Mecp2*^−/y^ mice experience worse sleep under conditions of elevated sleep pressure. This is particularly true of the differences in sleep amounts during the dark period.

Our findings demonstrate that the genetic loss of *MECP2* influences sleep architecture in a time-of-day-dependent manner. Previous studies have shown that circadian rhythms are affected in the absence of MECP2. *Mecp2*^−/y^ mice exhibit weakened locomotor activity rhythms, as well as a reduction in the number of neurons expressing vasoactive intestinal peptide (VIP) in the suprachiasmatic nucleus (SCN), the master clock located in the hypothalamus (Li et al., 2015). Sleep is regulated by two processes: the circadian clock (which determines the timing of sleep) and the sleep homeostat (which determines the need to sleep as a function of being awake) (Borbely, 1982). Our findings indicate that MECP2 may influence the circadian component of sleep regulation. In addition, although classical measures of sleep homeostasis are unaffected, the worsening of differences in sleep architecture relative to WT in response to SD suggests that the mutation alters how mice respond to sleep loss.

In summary, the loss of *Mecp2* in null mice significantly impacts sleep architecture in a time-of-day-dependent manner. *Mecp2*^−/y^ mice show increased NREM and decreased REM sleep during light periods but disrupted sleep patterns during dark periods that mirror the sleep-wake cycle disturbances observed in RTT patients. These findings establish a crucial link between MECP2 and circadian regulation of sleep, suggesting that while homeostatic sleep pressure mechanisms remain intact, the mutation alters how the brain responds to sleep loss by exacerbating baseline differences. Future research should explore the molecular pathways through which MECP2 influences circadian sleep regulation, particularly its interaction with the SCN, to develop targeted chronotherapeutic approaches for sleep disturbances in RTT patients. Additionally, investigating whether *MECP2*’s epigenetic regulatory functions directly affect clock gene expression could provide deeper insights into the mechanistic relationship between MECP2 and time-of-day regulation of sleep, potentially revealing new therapeutic targets for improving sleep quality in this vulnerable population.

## Credit authorship contribution statement

**Abrar Al Maghribi:** Conceptualization, Methodology, Software, Validation, Formal Analysis, Investigation, Data Curation, Writing - Original Draft, Writing-Review and Editing, Visualization. **Michael Rempe:** Software, Validation, Formal Analysis, Writing-Review & Editing, Visualization. **Elizabeth Medina:** Software, Investigation, Data Curation, Visualization, Writing-Review and Editing. **Kaitlyn Ford:** Investigation, Writing – Review & Editing. **Kristan Singletary:** Investigation, Writing – Review & Editing. **Caitlin Ottaway:** Visualization, Writing – Review & Editing. **Lucia Peixoto:** Conceptualization, Methodology, Validation, Resources, Data curation, Writing – review & editing, Supervision.

All authors approved the final manuscript as submitted and agree to be accountable for all aspects of the work. All authors have read and agreed to the published version of the manuscript.

## Data availability

All the data collected to support this study is available at https://staging.partners.org/sleepdata.org/datasets/mecp2zz and the code that was used to analyze the data is available at https://github.com/PeixotoLab/MeCP2Z.git

## Acknowledgements

We thank Dr. Huda Yahya Zoghbi for the valuable input using the Mecp2 mouse model to study Rett Syndrome our research.

## Appendix A. Supplementary data

Supplementary data available as additional files

Additional File 1

- Additional file 1 is a word document that includes all the supplementary figures (Additional Figure S1-S5) with their captions.

Additional File 2

- Additional file 2 is an excel sheet that includes details of all statistical data analysis that was applied in this study.

