## Additional File 1 for "Loss of MeCP2 leads to sleep deficits that are time-of-day dependent and worsen with sleep deprivation"

Supplementary Figure 1**. *Mecp2*$^{-/y}$ mice do not sustain sleep/wake states (REM Sleep Fragmentation) during baseline.** (A) Average number of bouts per hour of each arousal state (rows) during different portions of the day (columns). (B) Average duration of bouts. Wake top row, NREM sleep middle row, and REM sleep bottom row. Listed p-values indicate significant results of Two way-ANOVAs. The light period (hours 0-12), the first six hours of the dark period (hours 13-18), and the last six hours of the light period (hours 19-24) were tested separately. WT represented in black (n=10), *Mecp2*$^{-/y}$ represented in red (n=10).

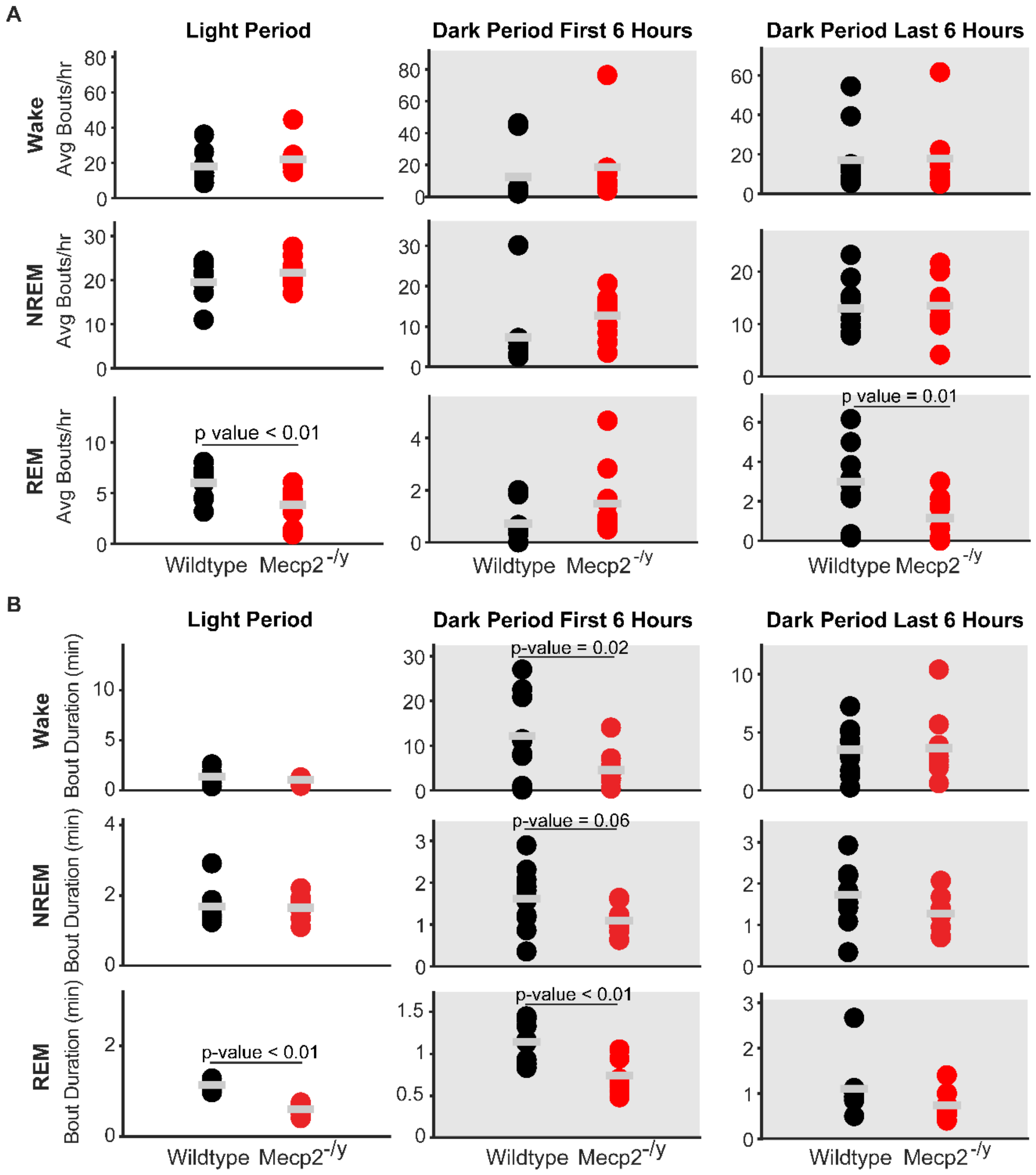

Supplementary Figure 2**. Baseline Spectra Power differences are similar in the light and dark periods.** Normalized EEG spectra power for the (A) light period (B) dark period at baseline (BL). In each case power was normalized to the mean power in all EEG frequencies averaged across all sleep states. The red line represents *Mecp2*$^{-/y}$ , and the black line represents the wild type. 95% confidence intervals are displayed around each spectrum, light gray for WT, and light red for *Mecp2*$^{-/y}$ . WT is represented in black (n=10), *Mecp2*$^{-/y}$ is

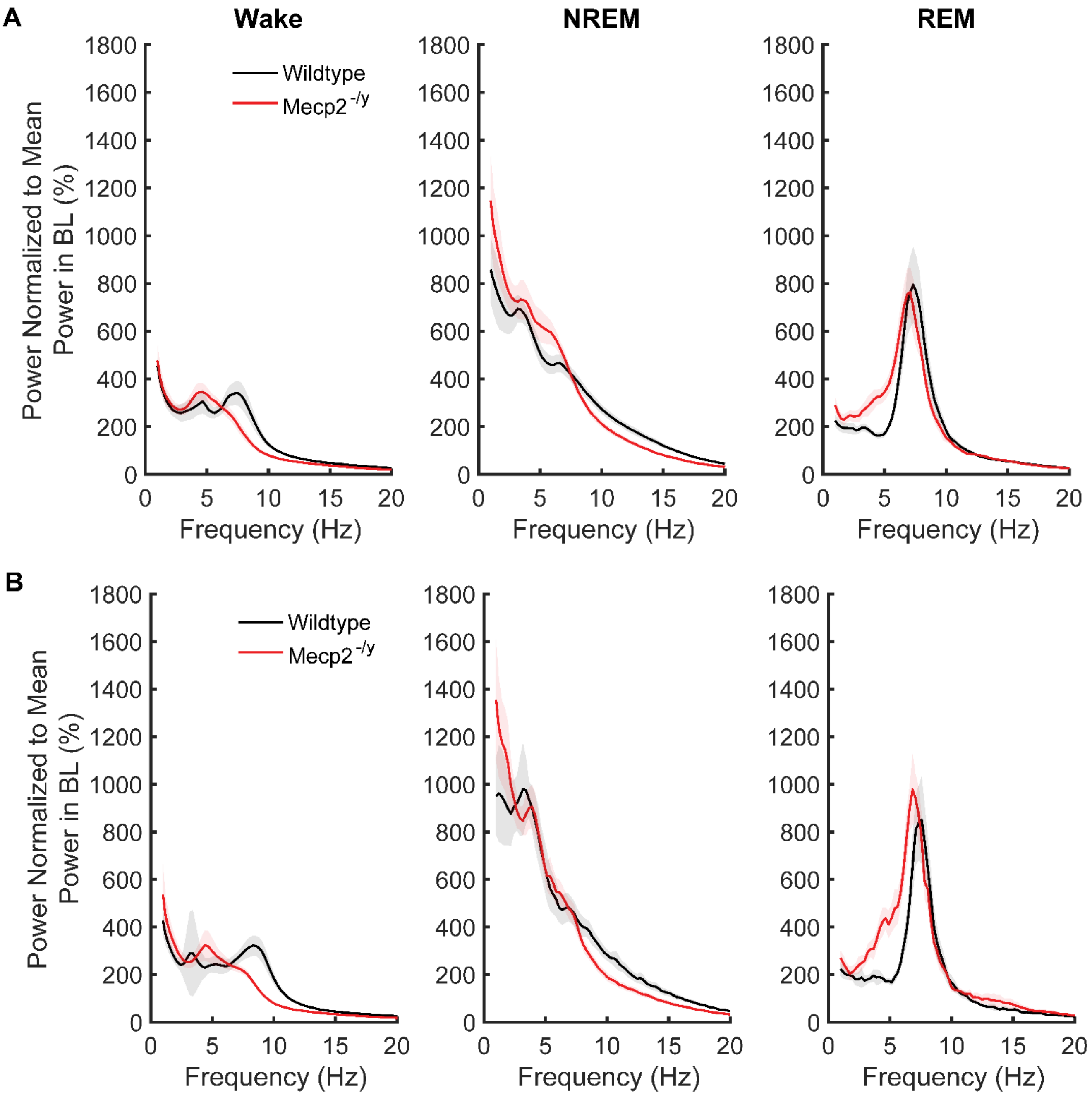

Supplementary Figure 3**. Sleep/Wake instability and worsened REM Sleep Fragmentation in *Mecp2*$^{-/y}$ Mice after Sleep Deprivation.** (A) Average number of bouts per hour of each arousal state (rows) during different portions of the day (columns). (B) Average duration of bouts. Wake top row, NREM sleep middle row, and REM sleep bottom row. Two way-ANOVA results. The light period (hours 0-12), the first six hours of the dark period (hours 13-18), and the last six hours of the light period (hours 19-24) were tested separately. WT represented in black (n=10), *Mecp2*$^{-/y}$ represented in red (n=10).

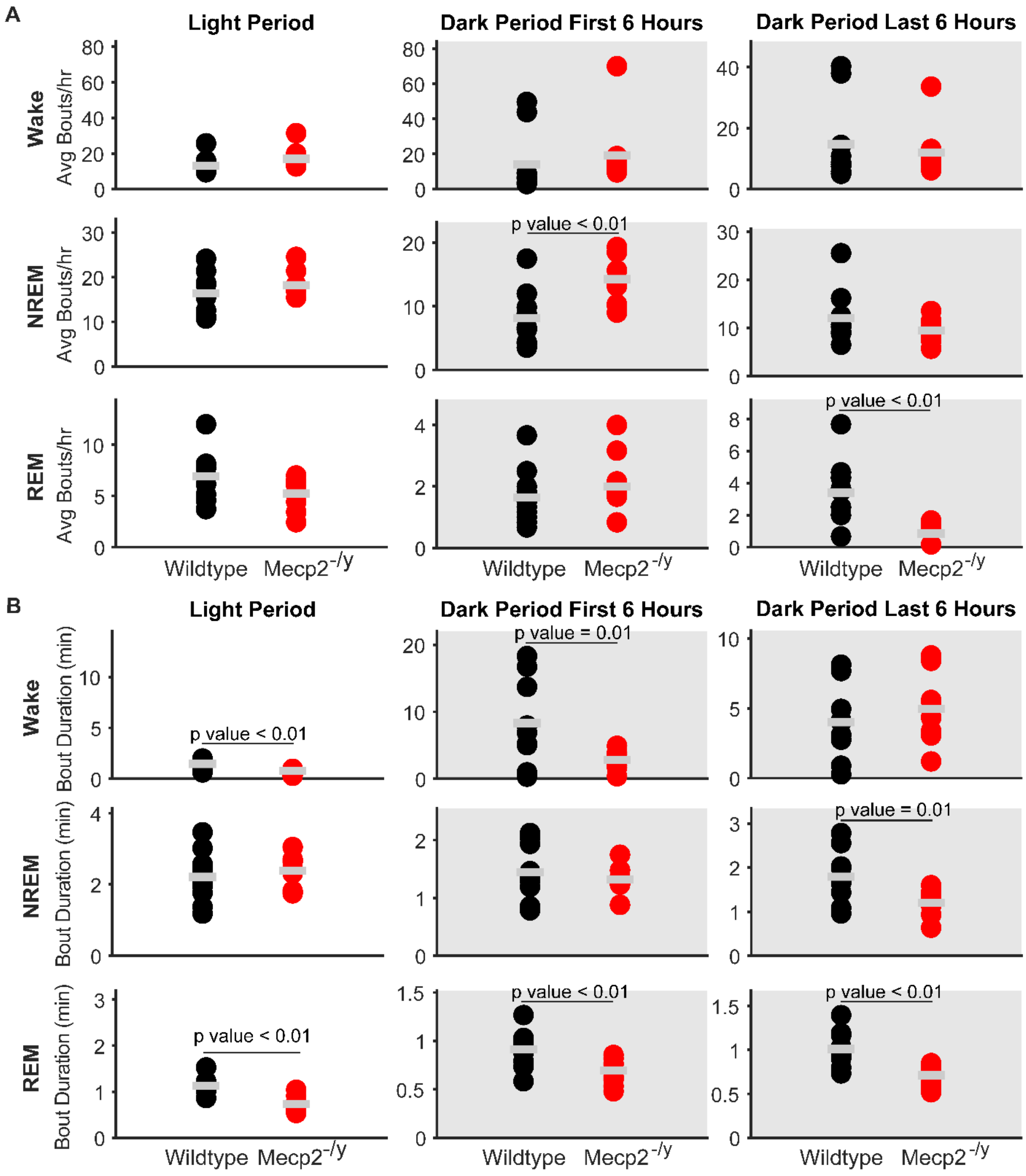

Supplementary Figure 4**. Recovery sleep spectral power.** Normalized EEG spectra power for the (A) light period (B) dark period of recovery day (C) over 24hrs post sleep deprivation. In each case power was normalized to the mean power in all EEG frequencies averaged across all sleep states. The red line represents *Mecp2*$^{-/y}$, and the black line represents the wild type. 95% confidence intervals are displayed around each spectrum, light gray for WT, and light red for *Mecp2*$^{-/y}$. WT is represented in black (n=10), *Mecp2*$^{-/y}$ is represented in red (n=10).

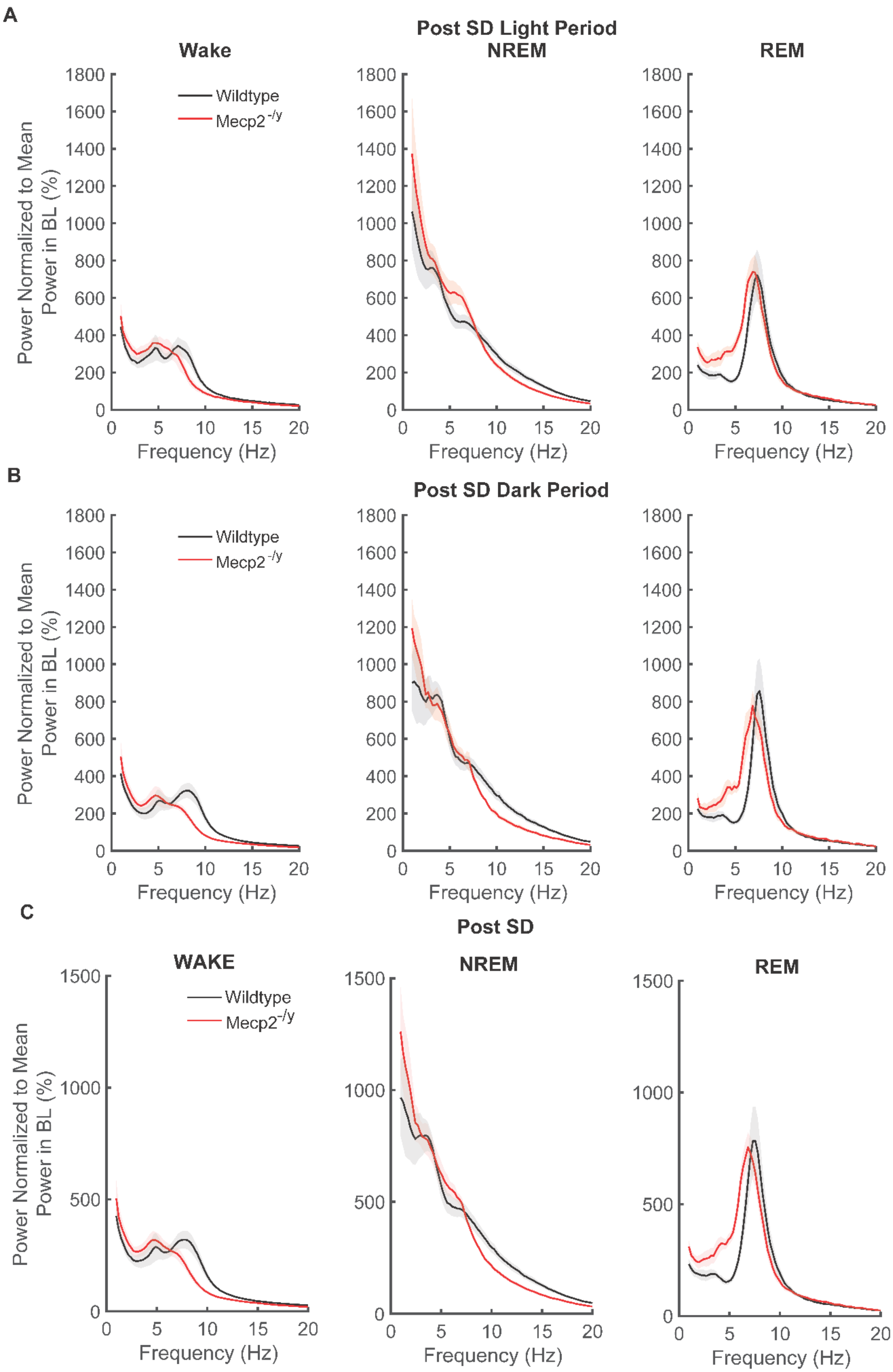
